# Flexible methods for estimating genetic distances from single nucleotide polymorphisms

**DOI:** 10.1101/004184

**Authors:** Simon Joly, David Bryant, Peter J. Lockhart

## Abstract

- With the increasing use of massively parallel sequencing approaches in evolutionary biology, the need for fast and accurate methods suitable to investigate genetic structure and evolutionary history are more important than ever. We propose new distance measures for estimating genetic distances between individuals when allelic variation, gene dosage and recombination could compromise standard approaches.
- We present four distance measures based on single nucleotide polymorphisms (SNP) and evaluate them against previously published measures using coalescent-based simulations. Simulations were used to test (i) whether the measures give unbiased and accurate distance estimates, (ii) whether they can accurately identify the genomic mixture of hybrid individuals and (iii) whether they give precise (low variance) estimates. The effect of rate variation among genes and recombination was also investigated.
- The results showed that the SNP-based GENPOFAD distance we propose appears to work well in the widest range of circumstances. It was the most accurate and precise method for estimating genetic distances and is also relatively good at estimating the genomic mixture of hybrid individuals.
- Our simulations provide benchmarks to compare the performance of different method that estimate genetic distances between organisms.

## Introduction

The last few decades have witnessed a methodological revolution in the field of population genetics. Model-based likelihood approaches have been propelled to the forefront of species and population level studies (e.g. Beaumont & Rannala 2004; Beaumont *et al.* 2002; Huelsenbeck *et al.* 2001). These changes have been made possible by the remarkable advances in computing technology and the application of computationally intensive Monte Carlo methodology. But even these sophisticated methods are facing critical challenges when confronted by the overwhelming quantity of data generated by massively parallel sequencing technologies. In many cases, state-of-the-art approaches in terms of models and methods cannot always accommodate population genomics data. Consequently, rapid methods that allow for investigations of patterns and processes still have their utility.

Our objective is to present new, flexible, and robust distance measures for estimating genetic distances from single nucleotide polymorphisms (SNPs) data. We focus on the estimation of distances between individuals (or organisms), even though the application of distances could certainly be useful in many other circumstances. There are good reasons to focus at the level of individuals rather than populations or species. Individuals are central to biology. Measurements based on morphology, spatial positioning, or genetics are generally performed at the individual level. Individuals are also the fundamental units of natural selection, the central concept of evolutionary biology. Finally, estimates of genetic relatedness between individuals can reveal correlations between genetic and phenotypic distances, spatial genetic structure across a landscape, species boundaries, and could be used for genetic or phylogenetic diversity surveys.

Although obtaining genetic distances among individuals seems relatively straightforward, there can be several complicating factors. One is the presence of allele variants in non-haploid individuals, which means that a single individual can be represented by two distinct sequences a given locus (Joly & Bruneau 2006; GökerPotts & Grimm 2008). Because of this, some obvious distance methods, such as taking the mean of all pairwise comparisons, could result in non-desirable results such as non null distance when an individual is compared with itself. Combining data from multiple loci also represents a challenge (Joly & Bruneau 2006). For instance, concatenation approaches are impossible as we cannot associate alleles from independent loci as they are segregating independently.

Polyploidy, which is defined by the presence of more than two genome copies in a nucleus, brings two other problematic issues: inheritance and gene dosage. Inheritance in polyploids is often unknown and could be either disomic or multisomic (Comai 2005). Polyploids are disomic if chromosomes group by pairs at meiosis, one example being homeologous chromosomes in allopolyploids. Polyploids are multisomic when chromosomes form multivalents. In many cases, inheritance of polyploid taxa is unknown or difficult to determine precisely. Some polyploids are even characterized by a mixture of inheritance modes. For instance, a marker could have mainly disomic inheritance with occasional multisomic inheritance, or different chromosomes could have different modes of inheritance within a genome (Wendel 2000).

Gene dosage is another problematic issue associated with polyploidy (Bruvo *et al.* 2004). In diploids, gene dosage is obvious: a homozygous individual has two identical alleles of the same gene and a heterozygous individual has two alleles with one copy of each. In polyploids, it is rare that we know the exact dosage of each allele in the genome. A tetraploid that has the observed nucleotide state ‘A’ at a position (i.e., it is homozygote) can only have genotype ‘AAAA’. However, a tetraploid individual with observed states ‘A’ and ‘T’ at a site could have the genotypes ‘ATTT’, ‘AATT’, or ‘AAAT’. The unknown dosage of these character states makes it more difficult to estimate precisely the genetic distances between polyploids. The situation can become even more complicated when there are more than two character states at a sequence site, a feature that becomes more likely in higher polyploids, or in comparisons involving individuals of different ploidy levels (Bruvo *et al.* 2004).

Finally, intragenic recombination can also complicate the estimation of genetic distances. With recombination, nucleotides within a marker can have different evolutionary histories. While this might be seen as beneficial since these regions provide independent although correlated outcomes of the evolutionary process, previous studies have shown that recombination could lead to an underestimation of the genetic distances (Schierup & Hein 2000; Bryant *et al.* 2003). So far, the impact of recombination in the context of individual genetic distances has not been properly investigated, though see Bryant *et al.* (2003).

Some methods have been proposed that can deal with some of these problems, but rarely with all of them (Joly & Bruneau 2006; Göker & Grimm 2008; Potts *et al.* 2014). Moreover, most of these methods use distances estimated by considering each marker as a whole, and are perhaps not well suited for the short length or SNP-based nature of the data obtained with massively parallel sequencing technologies (but see Potts *et al.* 2014). Finally, the performance of these methods has never been thoroughly tested using rigorous benchmarks.

Here, we propose four methods for estimating genetic distances between individuals from nucleotide sequence data. One of these is an adaptation of Nei’s genetic distance (Nei *et al.* 1983) for this specific problem, but the three other methods are novel. All methods are very general in that they can be applied to individuals of any ploidy level, but also when individuals of different ploidy levels are compared. We first describe the new methods and then compare them, and other previously published distances, using simulations. We finish by making recommendations on the use of distance measures in different contexts.

## Distance Definitions

We propose four new distance measures to calculate the genetic distance between individuals from sequence data. The main novelty of these measures is that they are all computed at the level of a nucleotide position in an alignment (an idea developed independently by Potts *et al.* 2014). We thus start by defining the distances at the nucleotide level, and explain later how they can be extended to strings of nucleotides, some potentially linked (within loci) and others unlinked (among loci). These measures assume that we know the nucleotides present at a given position in an individual but not necessarily gene dosage, which is typical for data obtained from genotyping or sequencing. All proposed distances are bounded between 0 and 1 and have the property that the distance between an individual and itself is 0.

### Matchstates

This measure looks at each nucleotide present at a given sequence site in one individual and checks if there is a nucleotide in the other individual that matches. More formally, consider a specific sequence site *i* that might be present in multiple copies in an individual. Note that in the present manuscript, gene copies are meant to designate copies of the same orthologous gene present in the individual or population. Let 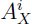 be the complete set of nucleotides for individual *X* at site *i* and let 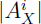 be the number of nucleotide states observed for individual *X* at site *i*. Let 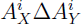 denote the set of nucleotides that belong to either 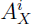 or 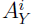, but not both. The matchstates distance between individual *X* and individual *Y* at site *i* is

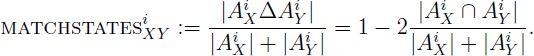

### genpofad

The genpofad measure is named after the pofad algorithm described by Joly and Bruneau (2006). The genpofad distance can be defined as one minus the ratio of the number of nucleotides shared between two individuals divided by the maximum number of nucleotides observed in either of the individuals at a given sequence site. Following the notation introduced above,

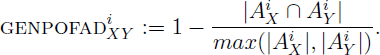

### mrca

The mrca distance measure (for Most Common Recent Ancestor) gives a distance of 0 whenever two individuals share at least one nucleotide at a given site and a distance of 1 otherwise. Formally, the mrca distance between individual *X* and individual *Y* is

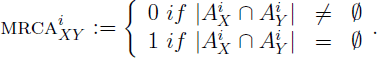

### nei

This distance is the application of Nei’s genetic distance (Nei *et al.* 1983) at the nucleotide level. The frequency of each nucleotide is estimated per site for each individual and then nei genetic distance between individual *X* and individual *Y* for site *i* is estimated as

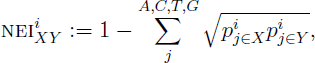

where 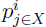 is the frequency of nucleotide *j* in individual *X* at site *i*. This formula is flexible as it can be easily applied among individuals from different ploidy levels. Gene dosage is assumed to be known, but it can also be used if it is unknown by giving equal weight to each nucleotide present.

### Extension to multiple sites and genes

The extension of all distance measures to many sites within a locus is easily done by taking the average distance over all DNA positions such as

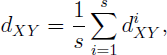

where *s* is the number of sites and 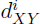 is the contribution of site *i* to the distance. If the mutations at each set are considered independent then an estimate of standard error is provided by the usual statistical formula

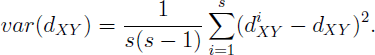

In some cases, it might be important to divide nucleotides into different loci, such as when several unlinked genes are sampled throughout the genome, each containing several linked nucleotides. We suggest distances be calculated first across sites within a marker to obtain distance matrices for each marker. Once this is done, one can compute a genome-wide distance matrix by taking the mean of all marker matrices. If the nucleotides cannot easily be divided into distinct loci, such as when we have a long contiguous sequence along a chromosome, the average distance over all DNA positions is appropriate because each site is then assumed to represent an independent assessment of the distance between the individuals.

### Implementation

These algorithms are all implemented in POFAD version 1.07 (Joly 2014, doi:10.5281/zenodo.11683). Note that the program can take either consensus sequences per individual using the IUPAC nucleotide ambiguity codes (Cornish-Bowden 1985) or it can take all sequences found in each individual if this information is available. The matchstates algorithm is also implemented in SplitsTree4 (Huson & Bryant 2006).

### Simulations

Computer simulations were performed to compare the performance of the distances in different situations. We evaluated three properties of the distance measures. First, we tested if the measures provide an unbiased and accurate estimate of distances between organisms. Second, we investigated how the different distances are able to detect the genomic mixture of hybrid individuals. Third, we evaluated how precise these different measures were.

We evaluated our new distance metrics together with those distance measures reviewed in Göker & Grimm (2008) which were relevant to the present context: the MIN distance and the Phylogenetic Bray-Curtis (pbc) distance (see Appendix for mathematical definitions). The frq and the entropy distance measures of Göker & Grimm (2008) were not investigated because they are not bounded between 0 and 1 and because they are more relevant in a context of host-parasite associations as originally described. Finally, we also evaluated the polymorphic *p* distance (Potts *et al.* 2014), hereafter pp, even if the distance is not bounded between 0 and 1 as it is similar to our proposed methods. The pp distance is defined using a nucleotide-based step matrix. Note that the pp distance used here differs slightly from that of the original publication; the distance obtained via the step matrix was divided by 2 following the suggestion of the authors (A. Potts, personal communication) so that the mutation from one non polymorphic nucleotide to another (e.g., A to G) receives a distance of 1 (and not 2 as in the original study).

### Accuracy of distance measures

To investigate whether the distance measures were accurate for estimating genetic distances between individuals, we simulated sequences in populations of tetraploid individuals (2*n*=4*x*) along a population tree using the coalescent, and estimated the genetic distances between individuals that have been evolving in independently evolving lineages for different periods of time. Gene sequences of 1000 bp were simulated using fastsimcoal2 vers. 2.5.0.2 (Excoffier & Foll 2011) on a population tree where populations had effective sizes of 5000 gene copies, which corresponds to 1250 tetraploid individuals. For the sake of simplicity, the effective population size used in the present manuscript is henceforth assumed to represent the number of gene copies in the population (*N_e(g)_*). The individuals compared belonged to populations that had been diverging independently for 0, 20000, 40000, 80000, 120000, and 200000 generations (*G*). A mutation rate of *μ* = 1^−7^ mutation per site per generation was used, which implies that *θ* = 2*N_e_(g)μ* = 0.001 in populations. Moreover, the divergence times scaled by the mutation rate (τ = *Gμ*) were equal to 0, 0.002, 0.004, 0.008, 0.012, and 0.02. These scaled divergence times (τ) are useful because they represent the expected number of mutations per site for a sequence from the divergence event in the population tree to the present time. However, the expected divergence times of the sequences is greater than the time of population divergence as the time to coalescence of the sequences in the ancestral population needs to be considered (Nei 1987; Edwards & Beerli 2000; Arbogast *et al.* 2002), which equals to the number of genes in the population (*N_e_(g)*) or *θ*/2 (Edwards & Beerli 2000). The expected genetic distance between two sequences is thus twice the coalescence time expectation, which is twice the time since the population divergence plus twice the expectation for the coalescent time in the ancestral population: d = 2μ + θ. Distance measures were compared to this expected sequence divergence and with the expected population divergence (2τ). Simulations were performed with two population sizes, *N_e_(g)*=5000 gene copies (*θ* = 0.001) and *N_e_(g)*=10000 (*θ*= 0.01), which were held constant throughout the tree. The larger population size increased the number of polymorphisms in individuals. All simulations were repeated 2000 times and each replicate consisted of one simulated DNA marker. Note that the scripts and code used to perform the simulations and data analysis have been deposited on Zenodo (doi: 10.5281/zenodo.12555).

### Impact of Rate Variation and Recombination

The impact of rate variation among genes was tested for the different methods. Rate variation was incorporated by multiplying the mutation rate by a random normal variate that had a mean of 1 and a standard deviation of 0.25. This implies that ca. 95% of the random variates falls between 0.5 and 1.5, which results in an overall 3-fold variation in rate between genes. These settings were selected because previous studies in plants have found that in general 90% of the genes encompass a 3-fold rate variation and that the distribution of rates is essentially normal (Senchina *et al.* 2003; Zhang *et al.* 2002), a pattern that seems very similar to that found in mammals (Hodgkinson & Eyre-Walker 2011). The same simulations as described previously for accuracy were performed with rate variation and they were compared with the results without rate variation for accuracy and precision (standard deviation of distances between replicates).

We also investigated the effect of adding recombination on the performance of the different distance measures. Recombination was included in the DNA markers at a rate of *r* = 2^−8^. The simulations were exactly identical to those described previously for distance accuracy, except that we simulated DNA sequence of both 1000 bp and 10000 bp. We compared the distributions of the results obtained with and without recombination using quantile-quantile plots.

### Estimation of the Genomic Mixture of Hybrids

To investigate how good the different distance measures are at detecting the genomic mixture of hybrid individuals, we estimated and compared the genetic distance of an allopolyploid individual with its two parents. For this, we simulated an allopolyploid speciation event. The parental species were tetraploids whereas the allopolyploid species was either octopolyploid with four gene copies coming from each parent or hexaploid with four copies coming from one parent and two from the other. This allowed us to test two ratios of parental genome contribution in the hybrid. Coalescent simulations were performed using multi labelled species trees (see Jones *et al.* 2013). This assumes that gene copies inherited from one parent are evolving independently from the gene copies inherited from the other parent in the allopolyploid, which is in concordance with a cytological definition of allopolyploidy. Consequently, the two parental copies in the allopolyploid can be simulated using two independent lineages for the allopolyploid species (Jones *et al.* 2013).

Gene sequences of 1000 bp were simulated with a mutation rate of *μ* = 1^−7^ on a population / species tree as described previously with a population size of *N_e_(g)* = 5000 genes copies (*θ* = 0.001). The divergence time between the parental species was fixed at 30000 generations (*G*), or τ = 0.003. Three different scenarios were investigated for the timing of the allopolyploid event: τ = 0 (in which case it is a first generation hybrid between the two parental species), τ = 0.001 or τ = 0.002. To investigate the hybrid mixture of the allopolyploid individual, we estimated a hybrid index that indicates the relative distance of the hybrid from its two parents:

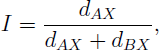

where *A* and *B* are the two parents and *X* the hybrid, and where *d_AX_* is the genetic distance between species A and the hybrid. The hybrid index (*I*) is bounded between 0 and 1 and an index of 0.5 indicates that the hybrid is equally distant to both parents. Cases where both *d_AX_* and *d_BX_* were equal to zero were given *I* = 0.5. All simulations were repeated 2000 times and each replicate consisted of one simulated DNA marker.

### Effect of the Number of Markers on Precision

We also estimated the impact of gene number on precision in the two previous simulation settings. For the precision of the genetic distance estimate, we used the simulations with *θ* = 0.001 and divergence time of τ = 0.012. For the hybrid index, we used the framework of the octopolyploid speciation event at τ = 0.001. In both cases, we evaluated the statistics (distance or hybrid index) estimated from 1, 2, 5, 10, 20, and 40 unlinked markers (taking the mean of all markers). Distances were estimated 500 times for each scenario and standard deviations among estimates were computed and plotted to investigate the decrease in standard deviation (i.e., increase in precision) with the number of markers for each method.

### Theoretical Considerations

Before comparing the different distance methods, it is relevant to note the similarities between the SNP-based methods proposed here and the previously published methods based on whole marker sequences. For example, MRCA is the same as MIN applied to a single nucleotide. As such, it is interesting to compare the performance of this pair of methods in the simulations. Moreover, the GENPOFAD distance is equivalent to the pofad algorithm of Joly & Bruneau (2006) when applied to a single nucleotide in diploid individuals. For a locus evolving under an infinite site mutation model without recombination, the GENPOFAD distance should give the same distance as pofad when extended to the whole locus. However, GENPOFAD has the advantage that it can be applied to individuals of any ploidy level, whereas pofad is limited to diploid individuals.

## Results

### Distance Accuracy

GENPOFAD and PP provided the most accurate estimates of the sequence divergence (2τ = *θ* (Fig 1), (Fig 2). However, in the simulations with small population sizes, both measures slightly underestimated the sequence divergence at high species divergence. Both genpofad and pp also provided an underestimated sequence divergence within populations (i.e., when τ = 0), suggesting that it is not a very accurate estimator of *θ*. Nevertheless, they were the best estimators of *θ* among the methods tested.

**Fig. 1.**
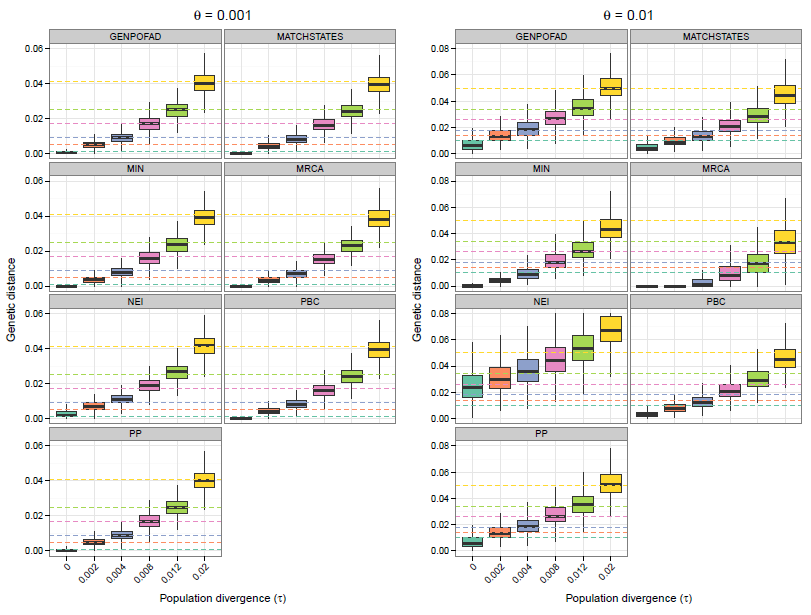
Boxplots showing the estimated divergence for several distance measures, compared to expected sequence divergence (d ; 2τ + *θ* dotted lines of the same colour as the boxes). Simulations were performed on a population tree with the coalescent using populations sizes of *θ* ; 0.001 (left panels) or *θ* ; 0.01 (right panels).

**Fig. 2.**
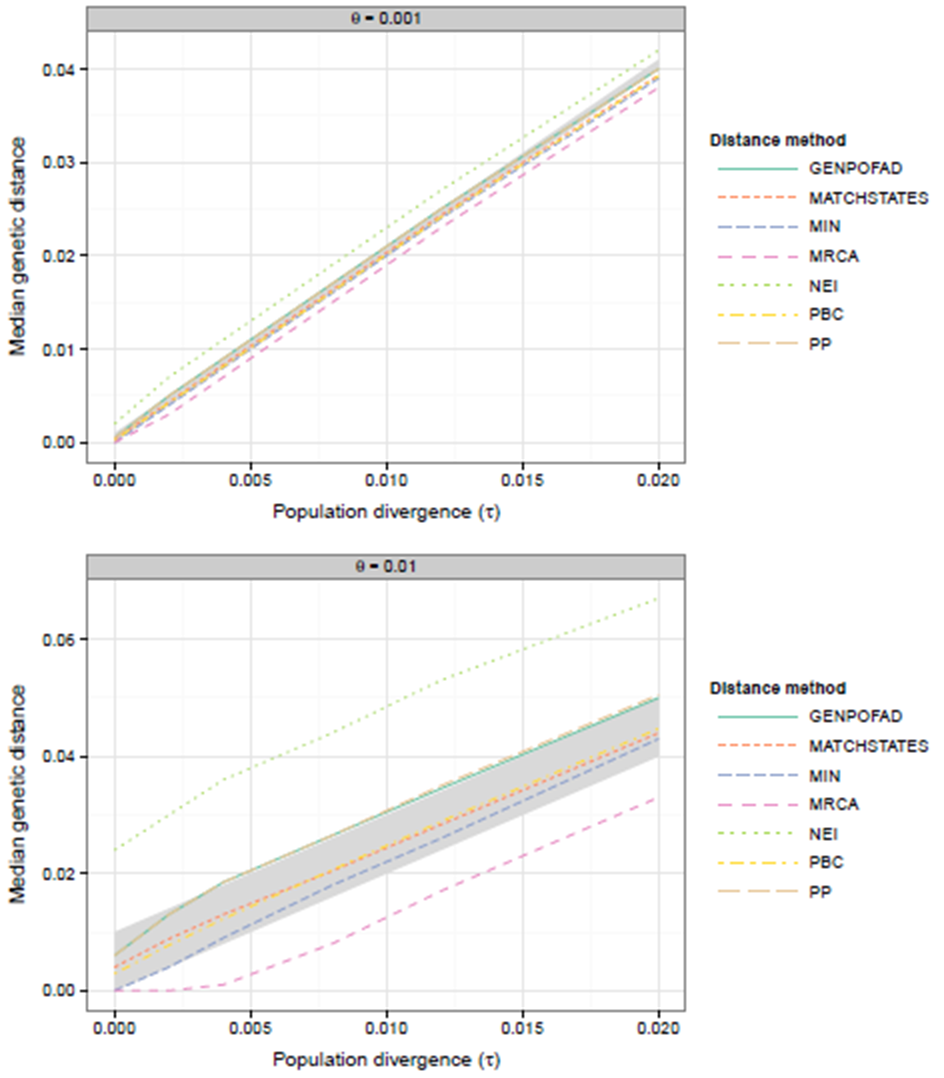
Plots showing the relationship between median estimated sequence divergence and population divergence for the distance methods, for two populations sizes. The gray area indicates the time range between the expected population divergence (d ; 2τ lower bound) and the expected sequence divergence (d ; 2τ + *θ* upper bound).

The other distances provided poor estimates of sequence divergence, although they sometimes had interesting properties. For instance, min provided an estimate closer to that of the population divergence time, even though this estimate gets biased with increasing divergence times (Fig 1), (Fig 2). Moreover, matchstates and pbc provided similar estimates that fell between the expected sequences divergence and the population divergence. The other measures either overestimated sequence divergence (nei) or underestimated population divergence (mrca) in all situations (Fig 1), (Fig 2).

### Rate Variation

Rate variation among genes did not affect the median distance in the simulations (Fig. S1), but it did result in a broader distribution of distances than when rate variation is absent (i.e., larger standard deviation; (Fig S1), (Fig 3). In general, all methods were similarly affected by the addition of rate variation, with the exception of mrca that seemed less affected than the other methods at large population sizes (Fig 3).

**Fig. 3.**
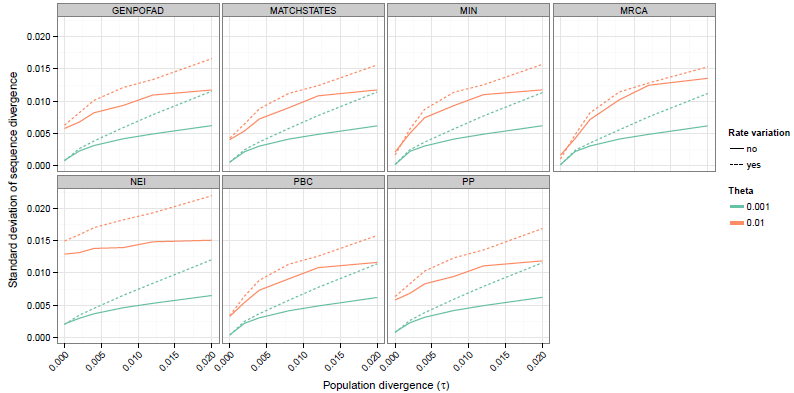
Plots illustrating the effect of rate variation among loci on the standard deviations of the distance estimates among replicates in a simulation. The solid and dashed lines give the results obtained without and with rate variation, respectively.

### Recombination

Overall, recombination did not have a strong impact on the results, and this was observed for all distance measures (Fig S2). To get a better appreciation of the impact of recombination, we compared the distributions of the distances obtained with and without recombination in simulations of 10000 bp sequences for different population divergences (Fig 4). The results obtained with longer sequences had the same trend as with shorter ones, but the effect of recombination is more evident (data not shown). Results showed that largest distances were smaller with recombination than without recombination. Similarly, recombination also resulted in fewer smaller distances, although this trend was not as constant amongst the scenario investigated. Interestingly, the bias on large distances becomes less important as population divergence (and thus sequence divergence) increases. All methods reacted similarly to the presence of hybridization, except perhaps for the nei distance which was less affected than others. Overall, recombination did not result in less accurate results than when recombination is absent. Instead, it resulted in smaller confidence intervals around the mean.

**Fig. 4.**
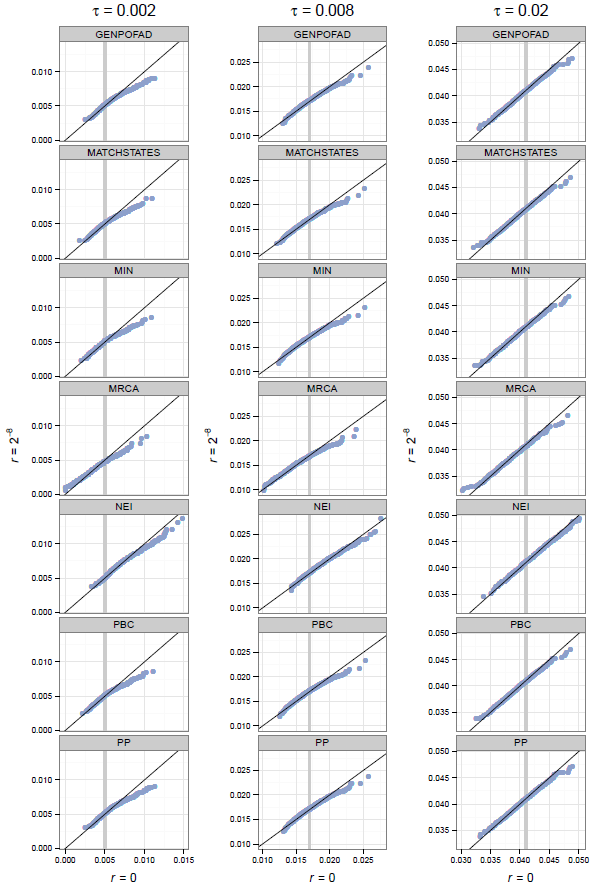
Quantile-quantile plots comparing the distance obtained for simulations performed at there population divergency, wuth recombination (y axis) and without recombination (x axis). The blank line gives expection for a perfect fit between the two distribution and gray lines give the expected sequence divergence (d ; 2τ + *θ*).Simulations were performed with DNA sequences of 10000 bp.

### Hybrid Genetic Mixture

Distance measures were evaluated as measures for estimating the contribution of parental genomes to allopolyploid hybrids. When the parents contributed an equal number of gene copies to the hybrid, all methods were accurate, but nei provided the most precise estimate of the hybrid index (Fig 5). genpofad and pp were the second best methods according to precision, followed very closely by pbc and matchstates. mrca and min provided imprecise estimates of hybrid index (Fig 5).

**Fig. 5.**
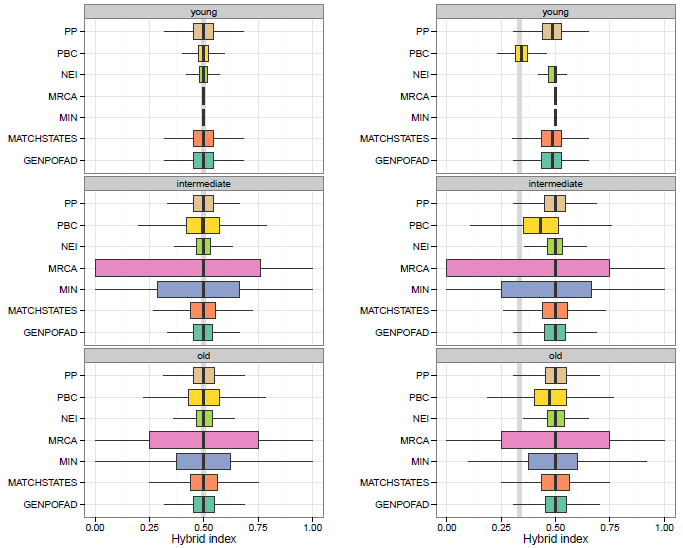
Boxplots showing a hybrid index (i.e., the relative contribution of each parental genome) for the different distance measures and for different times since the allopolyploid (hybridization) speciation event. The gray lines indicate the genomic mixtures that were simulated: one in which each parent contributed equally (1:1) to the allopolyploid (left panels) and another where one parent contributed twice the number of copies (2:1) than the other parent (right panels). The young, intermediate, and old events correspond to τ ; 0, τ ; 0.001, and τ ; 0.002, respectively.

No method provided an accurate hybrid index estimate when one parent contributed twice the number of gene copies than the other (Fig 5), but some methods performed better than others. pbc was by far the best method, followed by genpofad and matchstates. As before, mrca and min provided the worst estimates of the hybrid index. Also, if some evidence of an unequal contribution was visible for the young hybrid for genpofad and matchstates, evidence of unequal parental contribution for older hybrids was only observed with the pbc distance.

### Effect of the Number of Markers on Precision

Evaluation of the methods’ precision showed different results for the distance accuracy and for the hybrid index simulations. For the estimation of the genetic distance, all methods showed similar precision. Increase in precision (indicated by a decrease in the standard deviation among replicates) that accompanied the addition of more loci was similar for the different methods (Fig. 6a). The pattern was different for the precision of the hybrid index. The methods mrca and min were less precise than the others and they required more markers to converge on stable estimates (Fig. 6b). The remaining methods had a similar precision, although they could be ranked as followed for precision (from best to worst): NEI > genpofad = pp > matchstates > pbc (Fig. 6b).

**Fig. 6.**
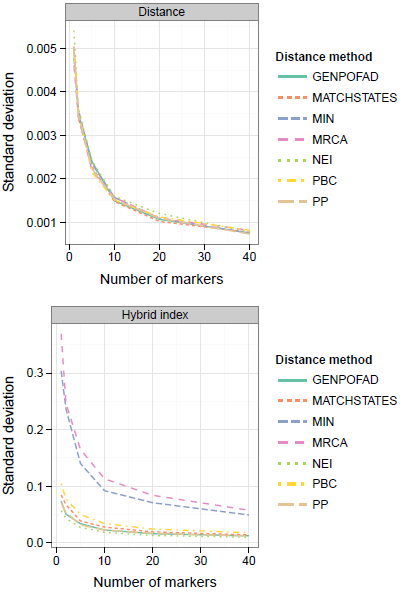
Plot showing the effect of the number of markers used on the precision of the genetic distances (upper panel) and the hybrid index estimates (lower panel). A small standard deviation indicates better precision. The simulation settings for the genetic distance were as for Figure 1a
Table Coding with a divergence time of 0.012 and for the hybrid index they were the same as those for Fig. 5a with an intermediate timing for the allopolyploid speciation events.

## Discussion

With the increasing use of massively parallel sequencing approaches in evolutionary biology, fast, accurate, and precise methods to investigate genetic structure and evolutionary history are required. Approaches based on concatenation are known to be inconsistent in some circumstances (Degnan & Rosenberg 2006; Salter Kubatko & Degnan 2007) and fully Bayesian approaches to population/species reconstruction (e.g. Heled & Drummond 2010; Liu *et al.* 2009) are computationally demanding with large numbers of markers. Even though faster coalescent alternatives for genomic studies are slowly being developed (Bryant *et al.* 2012), distance measures remain an important and useful tool, especially given the consistency properties of some indices (Liu *et al.* 2009; Mossel & Roch 2010).

Until now, the toolset of distance measures was limited for studying the relationships of individuals. Overcoming this shortcoming is critical given that individuals are the fundamental unit for many studies at the species level. The main problems encountered at this level are those of allelic variation and polyploidy. However, the potential presence of recombination in the nuclear genome and the SNP based nature of many contemporaneous studies represent further challenges. Thus we present here new distance measures that have the property that they are estimated at the nucleotide level in order to alleviate these biological complexities.

### Advantages of SNP-Based Distances

Interestingly, SNP-based distances did not suffer from the comparison with whole-sequence distances in our simulations. This is relevant because the simulation of long (1000 bp) sequences without recombination should advantage distances estimators based on whole sequences. To the contrary, the two most accurate methods for estimating genetic distances were SNP-based. This suggests that a SNP-based approach potentially offers more flexibility for estimating distances between organisms.

The presence of recombination did not seem to affect differentially the different methods as all reacted quite similarly. The only method that appeared less affected than the others was nei. Moreover, the observed underestimation of genetic distances in presence of recombination concurs with results from previous studies (Schierup & Hein 2000; Bryant *et al.* 2003). Finally, the observed reduced relative impact of recombination on the estimated distances at large divergence times can be explained by the fact that recombination can only occur within populations in our simulation settings and as such it cannot involves lineages that have been diverging for a long period of time.

Another important factor to consider is the length of markers. Massively parallel sequencing technologies generally result in short sequence markers. With such data, we expect that the relative advantage of distance measures based on the whole marker sequence to decrease with decreasing sequence length. Indeed, we can have an idea of that effect when going from 1000 bp sequences to SNP data by comparing the distances min and mrca as mrca is identical to min applied to a single SNP. Consequently, SNP-based methods are particularly well suited for SNP-based studies or for studies using short markers.

### Importance of Gene Dosage Information

Of the methods evaluated here, two can take into account exact gene dosage information: pbc and NEI. One would expect this type of information to be particularly important for estimating unequal genomic mixtures in hybrid individuals. This actually seems to be the case for pbc, which was the best method according to this criterion. However, nei did not appear to benefit from gene dosage information in the same situation. Our results tend to show, however, that gene dosage information is not critical for good performance in all situations. This is especially true for the estimation of genetic distances where the best methods did not use gene dosage information. This is a very encouraging result given that such information is rarely known precisely in genomic studies involving polyploids.

### Method Performances

In terms of genetic distance accuracy, the best methods were genpofad and pp, two SNP-based methods. At the population level, they both provided an underestimation of 9, even though they were better than all other methods in this aspect. The minimum allelic distance between individuals (min) provided an accurate estimate of the population / species divergence time at low sequence divergence. This observation concurs with previous studies that have shown this measure to be a consistent estimator of species distances in certain situations (Mossel & Roch 2010; DeGiorgio & Degnan 2014). But its utility in the present context appears limited given that it is biased with sequence divergence above approximately 0.005. Moreover, the simulations showed that this measure performs poorly when it comes to estimating the genomic mixture of hybrid individuals, both in terms of accuracy and precision. Interestingly, matchstates and the pbc provided estimates that fell between the expected sequence divergence and the species divergence, that is between 2τ + *θ* and 2τ.

Regarding hybrid mixture estimates, the best method was clearly pbc, which was the only method close to being accurate when estimating unequal contribution of the parents in the young age hybrid. Moreover, evidence for unequal contribution remained detectable for older hybrids, whereas that signal was lost for all other methods. Note that pbc used gene dosage information in the simulations, information that might not be always available in empirical datasets and that could affect its performance. Among other methods, genpofad and matchstates were slightly better as they showed slight evidence for the unequal parental contributions for the young hybrid and they provided precise estimates. The methods min and mrca were not precise and did not detect unequal parental contributions. This is not surprising as these methods essentially ignore polymorphisms by considering only the most similar nucleotides (mrca) or alleles (min).

## Recommendations

Any recommendations we make are necessarily based on the performance of the methods in the simulations used in this study and as such we cannot guarantee that the methods that performed well in the present study would do so in other circumstances. Nevertheless, in the light of our simulations, we believe that the best recommendation is to use the genpofad distance in general as this is the most accurate and precise method in terms of expected genetic distance and given that it is relatively good at estimating genomic mixture between individuals. Although pp had an almost identical behaviour to genpofad in our simulations, we prefer the use of genpofad for two reasons. First, because pp is based on a step matrix, it can theoretically allow distances higher than 1 at an alignment site if several nucleotide states are observed in one individual, an undesirable property for a distance. Second, its nature makes it more difficult to include gap characters as it would increase the complexity of the step matrix. In contrast, a gap character simply represents a fifth nucleotide state in the new methods proposed in this study.

In very specific situations, however, other methods might also be considered. For instance, in cases where population or species divergence times are of interest, the min distance may be considered at small genetic distances. Moreover, if gene dosage is known and genomic admixture is of main interest, then the pbc distance is probably the best choice. But most importantly, we hope that this study and the simulation framework we proposed for comparing the performance of genetic distance measures among organisms will stimulate the development and testing of further distances.

## Appendix

### Definition of Previously Published Distance Measures

In the following definitions based on whole markers sequences, A_X_ represents the complete set of alleles for individual *X* and |*A_X_*| is the number of alleles observed for individual *X*. Also, let d_ij_ be the genetic distance between alleles *i* and *j*.

### MIN Distance

The min distance was proposed by Göker & Grimm (2008) in the present context, but it had been often used in other contexts as well (e.g. Joly *et al.* 2009; Liu *et al.* 2009; Mossel & Roch 2010). It can be described as:

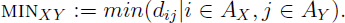

### Phylogenetic Bray-Curtis Distance (PBC)

The pbc distance was defined by Göker & Grimm (2008) as:

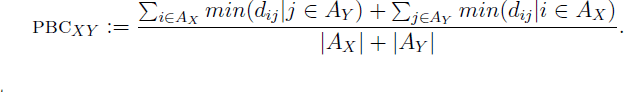

### PP Distance

The pp distance is a nucleotide-based distance (Potts *et al.* 2014). It estimates the distance between nucleotides using the step-matrix presented in Figure 1
Table Coding of Potts *et al.* (2014).

## Acknowledgements

The authors thank A. Potts and two anonymous reviewers for commenting on a previous version of the manuscript. This study was was supported by a NSERC Discovery Grant to SJ.

---

**Fig. S1.**
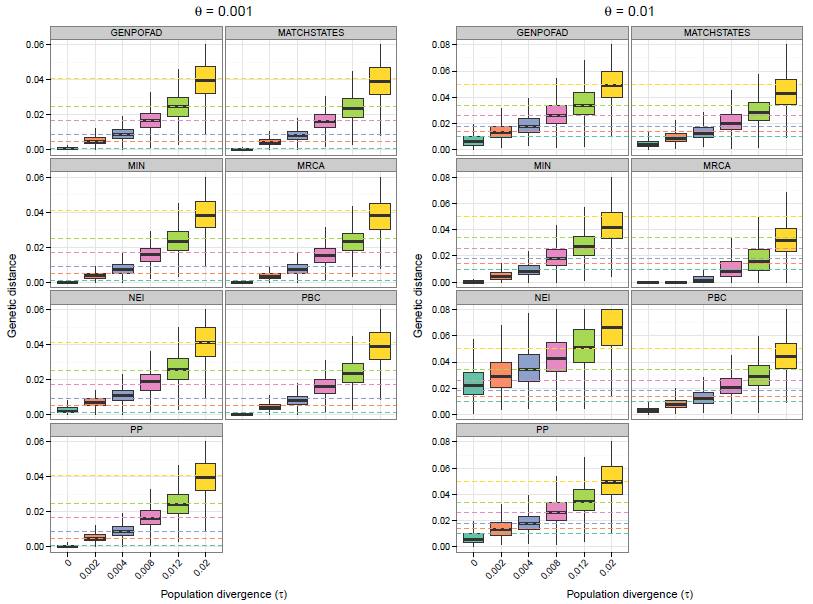
Boxplots showing the estimated sequence divergence for the distance measures in the presence of rate variation among genes, compared to expected sequence divergence (d ; 2τ + *θ*) dotted lines of the same colour as the boxes). Simulations were performed on a population tree with the coalescent using populations sizes of θ=0.001 (left panels) or 0.01 (right panels).

**Fig. S2.**
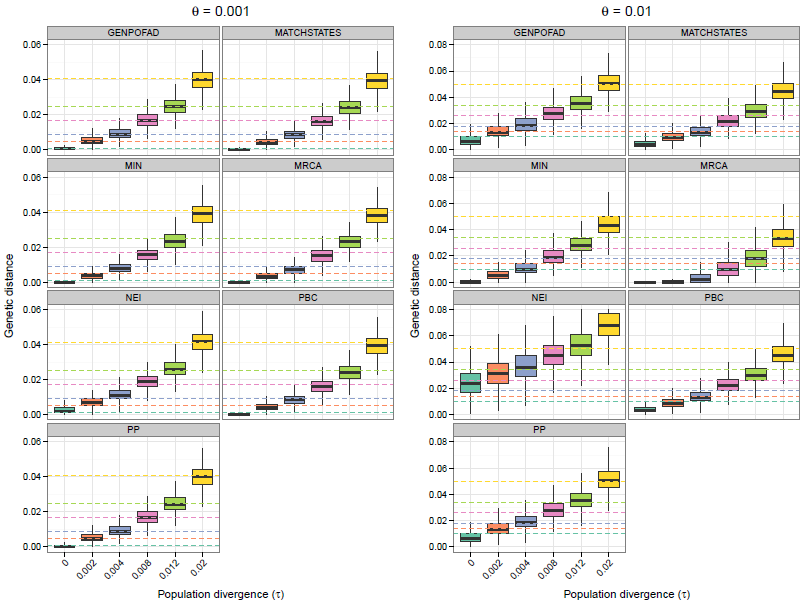
Boxplots showing the estimated sequence divergence for the distance measures in presence of recombination, compared to expected sequence divergence (d ; 2τ + *θ*) dotted lines of the same colour as the boxes). Simulations were performed on a population tree with the coalescent using populations sizes of *θ* ; 0.001 (left panels) or *θ* ; 0.01 (right panels). Simulations were performed with DNA sequences of 1000 bp.

